# Regulation of Na^+^,K^+^-ATPase isoforms and phospholemman (FXYD1) in skeletal muscle fibre types by exercise training and cold-water immersion in men

**DOI:** 10.1101/151035

**Authors:** Danny Christiansen, Robyn M. Murphy, James R. Broatch, Jens Bangsbo, Michael J. McKenna, David J. Bishop

**Author notes:** Corresponding Author: Robyn M. Murphy, Department of Biochemistry and Genetics, La Trobe Institute for Molecular Science, La Trobe University, Melbourne, Victoria, 3086, Australia.

## Abstract

Little is understood about the fibre type-dependent regulation of Na^+^,K^+^-ATPase (NKA) isoforms by exercise training in humans. The main aim of this study was therefore to assess the impact of a period of repeated exercise sessions on NKA-isoform protein abundance in different skeletal muscle fibre types in men. Post-exercise cold-water immersion (CWI) has been reported to increase oxidative stress, which may be one mechanism underlying increases in NKA-isoform expression. Thus, a second aim was to evaluate the effect of CWI on training-induced modulation of NKA-isoform abundance. Vastus lateralis muscle biopsies were obtained from nineteen men at rest before (Pre) and after (Post) six weeks of intense interval cycling, with training sessions followed by passive rest (CON, n=7) or CWI (10°C; COLD, n=5). Training increased (p<0.05) the abundance of NKAα_1_ and NKAβ_3_ in both type I and type II fibres, NKAβ_1_ in type II fibres, but was without effect on NKAα_2_ and NKAα_3_ (p>0.05). Furthermore, training decreased FXYD1 protein content in type I fibres, which abolished its fibre type-specific expression detected at Pre (p<0.05). CWI was without impact on the responses to training (p>0.05). These results highlight that NKA isoforms are regulated in a fibre type-dependent fashion in response to intense training in humans. This may partly explain the improvement in muscle Na^+^/K^+^ handling after a period of intense training. CWI may be performed without adversely or favourably affecting training-induced changes in NKA-isoform abundance.

**Summary in key points:**

- It is unclear how Na^+^,K^+^-ATPase (NKA) isoforms are regulated in different skeletal muscle fibre types by exercise training in humans, and the effect on phospholemman (FXYD1) protein abundance in different fibre types remains to be elucidated. We investigated the impact of six weeks of training on NKA-isoform protein abundance (α_1-3_, β_1-3_ and FXYD1) in type I and II muscle fibres in men.
- We show that intense interval training selectively increases the protein content of NKA α_1_ and β_3_ in both fibre types, β_1_ in type II fibres, and decreases FXYD1 in type I fibres.
- These results suggest the favourable impact of intense training on human muscle Na^+^/K^+^ regulation could be attributable, in part, to fibre type-dependent modulation of NKA-isoform abundance.
- Given that cold exposure has been shown to modulate cellular redox state, which has been linked to increased NKA expression, we also investigated the effect of exercise training plus cold-water immersion (CWI) on the fibre type-specific responses of NKA isoforms and FXYD1. We found that CWI was without effect on the responses to training.

**Abbreviations:** AMPKβ2, 5’ AMP-activated protein kinase subunit β_2_; CaMKII, Ca2+-calmodulin-dependent protein kinase isoform 2; COLD, cold-water immersion group; CON, control group; Ct, cycle threshold; CV, coefficient of variation; CWI, cold-water immersion; EDL, extensor digitorum longus; FXYD1, phospholemman isoform 1; HSP70, heat-shock protein 70; GXT, graded exercise test; K^+^, potassium; K_m_, Michaelis–Menten constant; MHC, myosin heavy chain; Na^+^, sodium; NF-1, neurofibromatosis type 1; NKA, Na^+^,K^+^-ATPase; ROS, reactive oxygen species; SDS-PAGE, sodium dodecyl sulphate polyacrylamide gel electrophoresis; SERCA1, sarco/endoplasmic reticulum Ca^2+^-ATPase isoform 1; VO_2peak,_ maximum oxygen uptake.

## Introduction

Skeletal muscle contractions invoke a cellular loss of potassium (K^+^) and a gain in sodium (Na^+^) ions, which has been associated with impaired muscle force development (McKenna *et al.*, 2008). These Na^+^ and K^+^ fluxes are counterbalanced, primarily, by one single, active transport system - the Na^+^,K^+^-ATPase (NKA) (Clausen, 2003). In skeletal muscle, the active NKA complex is composed of a catalytic α, a structural and regulatory β, and an accessory (phospholemman; FXYD) subunit (Clausen, 2013). Each of these subunits is expressed as different isoforms in both type I and II human skeletal muscle fibres (α_1-3_, β_1-3_ and FXYD1) (Thomassen *et al.*, 2013; Wyckelsma *et al.*, 2015). It has been shown in rats that differences in NKA-isoform abundance between these fibre types could be responsible for the discrepancy between muscles in their capacity for Na^+^ and K^+^ transport (Kristensen & Juel, 2010a). Thus, fibre-type specific study of NKA adaptation is fundamentally important to understand the regulation of Na^+^/K^+^ homeostasis and muscle function.

One study has investigated training-induced effects on NKA-isoform abundance in isolated type I and II muscle fibres from humans. In that study, Wyckelsma *et al.* (2015) found no effect of four weeks of repeated-sprint training (12 sessions of three sets of 5 × 4-s running sprints with 20 s of rest between sprints and 4.5 min of rest between sets) on α-isoform abundance (α_1_, α_2_ and α_3_). In contrast, the abundance of these isoforms has been reported to change in whole-muscle samples in response to a substantially (~12-18 times) greater training volume (Mohr *et al.*, 2007; Iaia *et al.*, 2008; Bangsbo *et al.*, 2009). Thus, the chosen volume and duration of training may have reduced the likelihood of detecting changes in NKA with training in the study by Wyckelsma *et al.* (2015). It is by no means clear how the NKA isoforms are regulated by exercise training in the different fibre types in humans, and the effect of exercise training on FXYD1 abundance in different muscle fibre types remains to be elucidated.

There is great interest in the use of cold-water immersion (CWI) to optimise muscle recovery, and how it may affect adaptations to exercise training (Versey *et al.*, 2013; Broatch *et al.*, 2014; Roberts *et al.*, 2015). But no study has investigated the effect of post-exercise CWI on muscle NKA protein content. In humans, cold exposure has been shown to increase the systemic level of norepinephrine (Gregson *et al.*, 2013), which has been linked to greater oxidative stress (Juel *et al.*, 2015). In many cell types, cold exposure has also been reported to perturb redox homeostasis and increase the production of reactive oxygen species (ROS) (Selman *et al.*, 2002). Given these cellular stressors could be important for the isoform-dependent modulation of NKA-isoform expression in response to exercise in humans (Murphy *et al.*, 2008), post-exercise CWI may be a potent stimulus to affect NKA-isoform content in humans. However, this hypothesis remains currently untested.

The aims of the present study were to evaluate the effect of exercise training on NKA-isoform protein abundance (α_1-3_, β_1-3_ and FXYD1) in type I and type II skeletal muscle fibres in humans and to examine if these adaptations could be modified by post-exercise CWI. In addition, as limited information is available on the reliability of western blotting for NKA isoforms, and to support our interpretation of data, we determined the technical error of this method for each of the NKA isoforms.

## Methods

### Ethical Approval

This study was approved by the Human Research Ethics Committee of Victoria University, Australia (HRE12-335) and conformed to the Declaration of Helsinki. The participants received a detailed oral and written, plain-language explanation of the procedures, potential risks, and benefits associated with the study before providing oral and written consent.

### Participants

Nineteen healthy males volunteered to participate in this study. Their age, body mass, height and peak oxygen uptake (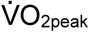) were (mean ± SD) 24 ± 6 y, 79.5 ± 10.8 kg, 180.5 ± 10.0 cm and 44.6 ± 5.8 mL·kg^-1^·min^-1^, respectively. The participants were non-smokers and engaged in physical activity several days per week, but were neither specifically nor highly trained.

### Experimental design

Before the first biopsy session, participants reported to the laboratory on two separate occasions to be accustomed to the exercise protocol and recovery treatments. During this period, they also performed a graded exercise test (GXT) to volitional exhaustion on an electromagnetically-braked cycle ergometer (Lode, Groningen, The Netherlands) and their 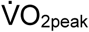 was assessed in accordance with published methods (Broatch *et al.*, 2014). These visits were concluded at least 3 days prior to the first biopsy session and separated by a minimum of 24 h. We utilised a parallel, two-group, longitudinal study design (Fig. 1). Participants were matched on their pre-determined 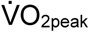 and randomly assigned by a random-number generator (Microsoft Excel, MS Office 2013), in a counter-balanced fashion, to one of two recovery treatments: Cold-water immersion (COLD, *n =* 9) or non-immersion rest at room temperature (CON, *n =* 10). The assigned recovery protocol was performed upon completion of every intense, sprint interval exercise session during the six weeks of training. A muscle biopsy was obtained at rest before the first, (Pre) and approximately 48-72 h after the last (Post), training session. These samples were used to determine protein abundance in type I and II muscle fibres. This study was part of a larger research project investigating the effects of post-exercise CWI on muscle adaptation in humans. Due to insufficient size of some muscle biopsies allocated to this part of the study, protein abundance was determined in twelve of these participants (CON, *n =* 7; COLD, *n =* 5).

**Figure 1.**
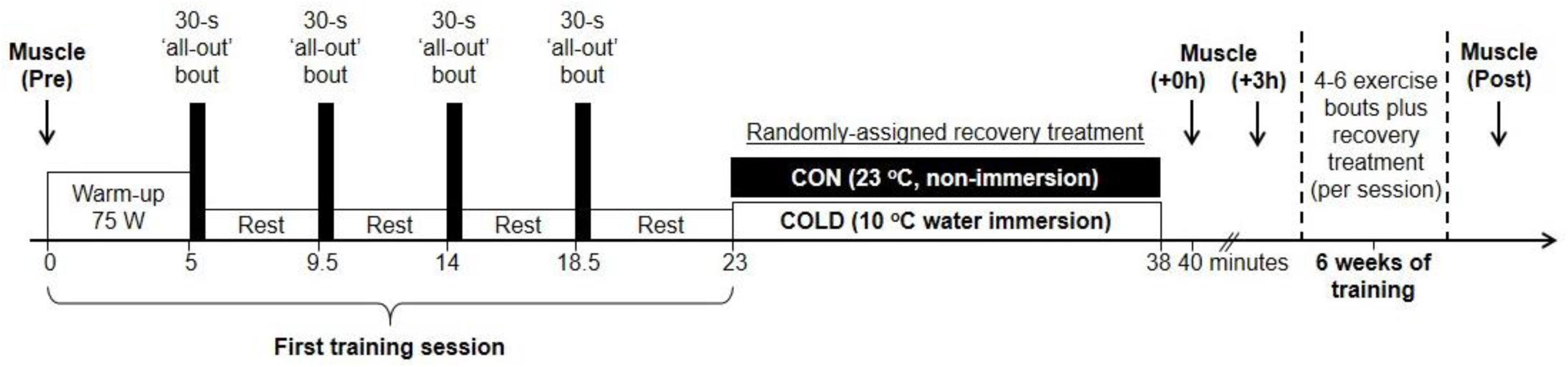
A time-aligned schematic representation of the experimental setup. Muscle was sampled at rest before exercise (Pre), 2 min post (+0h) and 3 h after (+3h) 15-min of passive rest at room temperature (CON group) or cold-water immersion up to the umbilicus (~10°C; COLD group) that followed the first intermittent, sprint training session. A final biopsy was sampled at rest 48-72 h following the last training session (Post) of six weeks of intense interval training either without (CON) or with cold-water immersion (COLD) after each training session. This study was part of a larger study. Muscle sampled at Pre and Post only was used for the current analyses.

### Exercise protocol and training

All experimental and training sessions took place in the Exercise Physiology Laboratory at the Institute of Sport, Exercise and Active Living (ISEAL), Victoria University (Melbourne, Australia). The participants completed the first training session on the same electrically-braked cycle ergometer as used during the familiarisation. After a 5-min warm-up at a constant absolute intensity (75 W), the participants performed four 30-s maximal-intensity (‘all-out’) sprint efforts at a constant relative flywheel resistance of 7.5 % of body mass, interspersed by 4 min of passive recovery in which they remained seated with their legs resting in the pedals. Each effort was commenced from a flying start at ~120 rpm, brought about by the investigators’ manual acceleration of the flywheel before each effort. The participants remained seated in the saddle throughout the entire session. Augmented verbal feedback was provided to the participants by one investigator in a consistent manner throughout each effort. This protocol was repeated in every training session and was performed under standard laboratory conditions (~23°C, ~35% relative humidity). To ensure a progressive physiological stimulus, participants performed four sprint repetitions in weeks 1-2, five in weeks 3-4, and six in weeks 5-6. Pedal resistance was modified (7.5-9.5% of body mass) during training to ensure a minimum fatigue-induced decline in mean power output of 20 W·s^-1^.

### Recovery treatments

Five minutes after termination of the sprint interval exercise, participants commenced their designated 15-min recovery treatment, consisting of either rest in a seated posture with the legs fully extended on a laboratory bed at room temperature (~23°C, CON) or 10°C water immersion up to the umbilicus in the same position in an inflatable bath (COLD; iBody, iCool Sport, Miami QLD, Australia). The water temperature was held constant by a cooling unit (Dual Temp Unit, iCool Sport, Miami QLD, Australia) with constant agitation.

### Muscle sampling

Muscle was sampled from the *vastus lateralis* muscle of the participants’ right leg using a 5-mm Bergström needle with suction. In preparation, a small incision was made at the muscle belly through the skin, subcutaneous tissue and fascia under local anaesthesia (5 ml, 1% Xylocaine). Separate incisions were made for each biopsy and separated by approximately 1-2 cm to help minimise interference of prior muscle sampling on the physiological response. The participants rested on a laboratory bed during each sampling procedure and the biopsies were obtained after ~30 min of rest in the supine position. Immediately after sampling, samples were rapidly blotted on filter paper to remove excessive blood, and instantly frozen in liquid nitrogen. The samples were then stored at - 80°C until subsequent analyses. The incisions were covered with sterile Band-Aid strips and a waterproof Tegaderm film dressing (3M, North Ryde, Australia).

### Dissection of single-fibre segments

Skeletal muscle single-fibre segments were collected and prepared for western blotting as previously described (Murphy, 2011). Approximately 105 (range: 17-218) mg w.w. of muscle was freeze dried for 28 h, providing 24 (3.4-49.0) mg d.w. muscle for dissection of individual fibres. By use of fine jeweller’s forceps, a minimum of 20 single-fibre segments were separated from each biopsy sample in a petri dish under a light microscope at room temperature (~45-60 min per biopsy). A camera (Moticam 2500, Motic Microscopes) attached to a monitor was connected to the microscope to manually measure, by use of a ruler, the length of each fibre segment. The mean (range) segment length was 1.5 (1.0-3.2) mm and a total of 520 fibre segments were collected. After dissection, segments were placed in individual microfuge tubes with the use of forceps and incubated for 1 h at room temperature in 10 μL SDS buffer (0.125 M Tris-HCl, 10% glycerol, 4% SDS, 4 M urea, 10% mercaptoethanol, and 0.001% bromophenol blue, pH 6.8), after which they were stored at −20 °C until analysed.

### Preparation of muscle fibre pools

One half of each solubilised fibre segment was used to qualitatively determine fibre type by western blotting with antibodies against MHC I and II (see section on *Immunoblotting*). The other half of each fibre segment was grouped with other fibre segments from the same biopsy according to MHC expression, to form samples of type I or II fibres from each biopsy, similar to the procedure described previously (Kristensen *et al.*, 2015). The mean ± SD number of fibre segments analysed per participant was 36 ± 3 (427 segments in total). The number of fibre segments included in each pool of fibres per biopsy before and after training, respectively, was 8 (range 4-14) and 5 (range 2-10) for pools of type I, and 10 (range 6-13) and 12 (range 7-16) for pools of type II, fibres. Hybrid fibres (i.e., expressing multiple MHCI and IIa isoforms) were excluded from analysis (*n* = 22, ~4 %). Type IIx fibres, classified by a lack of MHC I and II content despite protein present in the sample, were also excluded (*n =* 5, < 1%). Some lanes on the Stain Free gel were empty, indicating that no fibre was successfully transferred into the microfuge tube (*n =* 8, ~1.5 %). Based on agreement between two independent researchers, who conducted visual inspections of the blots, fibre segments and points on the calibration curves were excluded from analysis if their band was unable to be validly quantified due to noise on the image caused by artefacts or if they were too faint or were saturated.

### Calibration curves

To be able to compare across gels, and to ensure that blot density was within a linear range of detection (Murphy & Lamb, 2013), a four-point calibration curve of whole-muscle crude homogenate with a known amount of protein was loaded onto every gel. The homogenate was prepared from an equal number (*n* = 5) of pre- and post-training freeze-dried, fibre-type heterogeneous muscle samples. The samples were manually powdered using a Teflon pestle in an Eppendorf tube and then incubated for 1 h at room temperature in SDS buffer (0.125 M Tris-HCl, 10% glycerol, 4% SDS, 4 M urea, 10% mercaptoethanol, and 0.001% bromophenol blue, pH 6.8). The protein concentration of the homogenate was estimated by Stain Free gel electrophoresis (Bio-Rad, Hercules, CA). The intensity of the protein bands was compared to a standard curve of mixed human muscle homogenate with a known protein concentration.

### Immunoblotting

To determine MHC isoform abundance in each fibre segment, half the solubilised segment (5 μL) was loaded onto 26-wells, 4-15% or 4-20% Criterion TGX Stain Free gels (Bio-Rad Laboratories, Hercules, CA). Each gel was loaded with 20 segments from the same participant (*n* = 10 for pre and post training), two protein ladders (PageRuler, Thermo Fisher Scientific) and a calibration curve. NKA α_1-3_, β_1-3_, and FXYD1 protein in the muscle fibre pools was quantified by loading 15 μg w.w. muscle per sample, along with a calibration curve, onto the same gel using the same gel type as per above. Pools of type I and II fibres from Pre and Post from the same participant were loaded onto the same gel. Gel electrophoresis was performed at 200 V for 45 min. After UV activation for 5 min on a Criterion Stain Free Imager (Bio-Rad), proteins in gels were wet-transferred to 0.45 μm nitrocellulose membrane for 30 min at 100 V in a circulating, ice-cooled bath using transfer buffer (25 mM Tris, 190 mM glycine and 20% methanol). The current was on average ~0.50-0.75 mA and did not exceed 0.95 mA. After transfer, membranes were incubated for 10 min in Pierce Miser solution (Pierce, Rockford, IL, USA), washed five times in double-distilled H_2_O, and blocked for 1.5 h in blocking buffer (5% non-fat milk in Tris-buffered saline Tween, TBST) at room temperature with rocking. Membranes containing fibre pools were cut horizontally at 170 kDa, 70 kDa and 25 kDa to re-determine fibre type (MHC isoforms, ~200 kDa) and to quantify one NKA α isoform (~100 kDa), along with one β isoform (~50 kDa) and FXYD1 (~12 kDa) on the same membrane. Membranes were incubated with rocking overnight at 4°C (preceded or followed by 2 h at room temperature) in primary antibody diluted in 1% bovine serum albumin (BSA) in phosphate-buffered saline with 0.025% Tween (PBST) and 0.02% NaN_3_ at concentrations detailed in Table 3. To improve the visualisation of fibre type, the top portions of each membrane (>170 kDa) were stripped between MHCIIa (1^st^) and MHCI (2^nd^) probes for 30 min at 37 °C in western blot stripping buffer (#21059, Thermo Fisher Scientific, MA USA). NKA isoforms were quantified as first probes, except for α_2_ and β_3_, which were quantified as second probes following quantification of α_1_ and β_1_, respectively. Use of different host species for α_1_ (mouse) and α_2_ (rabbit), and the distinct molecular bands of β_1_ and β_3_ allowed their quantitative assessment on the same membrane. No stripping of these membrane portions were performed. After incubation in primary antibody, membranes were washed three times in TBST and incubated for 1 h at room temperature in horseradish peroxidase (HRP)-conjugated secondary antibody (goat anti-mouse immunoglobulins or goat anti-rabbit immunoglobulins; Pierce, Rockford, IL, USA) diluted 1:20.000 with 5 % non-fat milk. After another three membrane washes in TBST, protein bands were visualised using enhanced chemiluminescence (SuperSignal West Femto, Rockford, Pierce, IL, USA) on a ChemiDoc MP imaging system (Bio-Rad). Quantification of bands was performed in Image Lab 5.2.1 (Bio-Rad). Linearity between blot signal (density) and tissue loaded for calibration curves was established on every membrane. The same researcher was responsible for performing all western blots included in this study.

### Antibodies for immunoblotting

Full details for the primary antibodies are shown in Table 1. Validation of antibodies is shown in Fig. 2 and was performed with positive and negative controls using mouse (EDL, soleus and brain), rat (EDL, soleus, brain, kidney and cardiac muscles) and human (breast cancer cell lines, embryonic kidney cells and skeletal muscle) tissues. The rats (Sprague Dawley, six months old) and mice were sedentary and healthy. Rat and mouse tissue was obtained from animals used under La Trobe University Animal Ethics Committee (approval AEC 14-33). Antibodies used to detect myosin heavy chain slow- (type I, #A4.840) and fast-type (type IIa, #A4.74) isoforms were produced using the entire immunogen sequence (MYH7 and MYH2, respectively). The former antibody recognises a C-terminus epitope, whilst the latter remains to be mapped (DSHB).

**Table 1.**
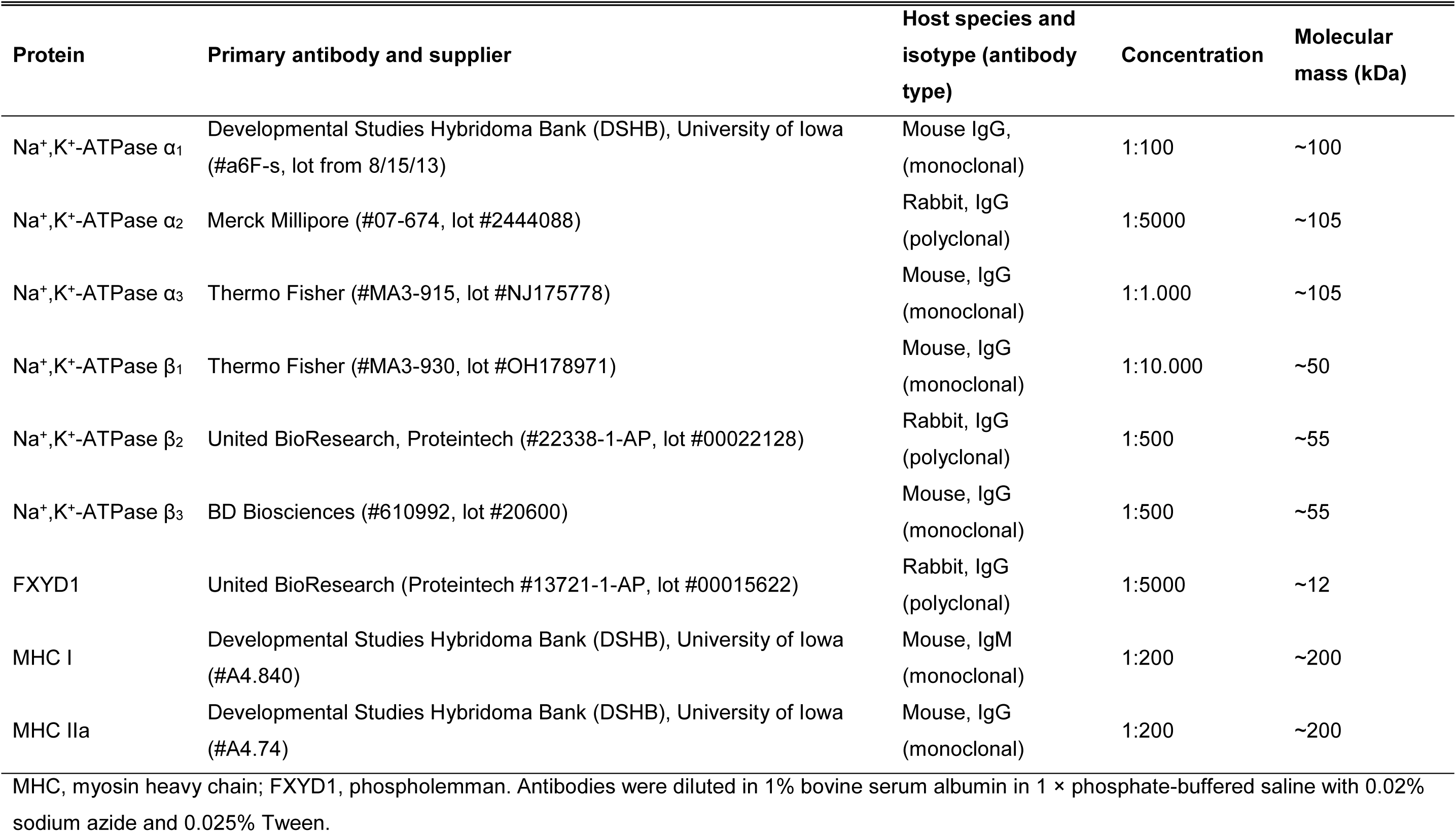
Primary antibodies used for quantification of protein abundance of Na^+^,K^+^-ATPase isoforms, phospholemman (FXYD1) and myosin heavy chain isoforms in groups of type I or type II human skeletal muscle fibres

**Figure 2.**
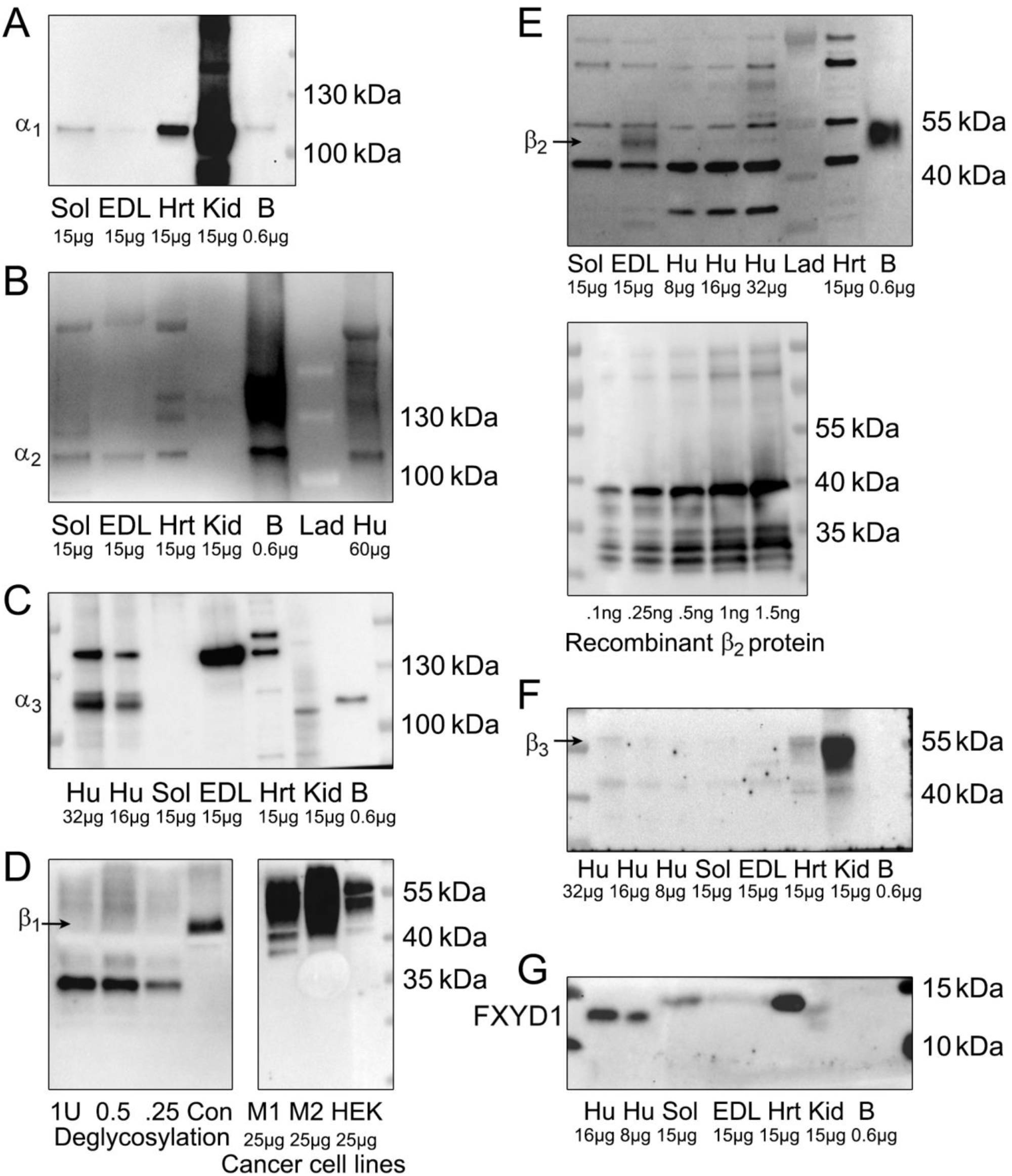
Validation of antibodies used to quantify Na^+^,K^+^-ATPase (NKA) isoforms and FXYD1. Crude samples of human vastus lateralis (Hu) and rat skeletal muscles (EDL, extensor digitorum longus; Sol, soleus), rat cardac muscle/heart (Hrt), kidney (Kid) and brain (B), breast cancer cell lines (M1, MDA-MB-231; M2, MCF10.Ca1d) and a control cell line (HEK, human embryonic kidney 293) were loaded onto 4-15% gradient, Criterion Stain Free gels. After SDS-PAGE, proteins were wet-transferred onto 0.45 μm nitrocellulose membrane. Membranes were incubated with antibodies raised against each of the NKA isoforms (α_1-3_ in A, B and C; β_1-3_ in D, E and F, respectively) or FXYD1 (G), post-treated with specific secondary antibodies and imaged using chemiluminescence. Isoform bands and molecular weight markers (lad) are shown in each image. In D, deglycosylation of human skeletal muscle samples were performed using PNGase incubation (3 h) at concentrations indicated in units (U). Recombinant β2 protein (#Ag17818, Protein Tech) identical to the 64-171 aa derived from E.coli, PGEX-4T with N-terminal GST was used to verify the specificity of the β_2_ antibody (E, *bottom panel*). See *Methods* section for additional details.

### Reliability of western blotting

The reproducibility of western blotting for each of the NKA isoforms and FXYD1 was determined from triplicate western blots for each of the proteins and is expressed as the coefficient of variation (CV; Fig. 3C). The calibration curves from whole-muscle, crude homogenate, which were loaded on every gel, were used for the analysis. Protein abundance and total protein for each amount of protein loaded was determined by normalising the density for a given loading amount to that of the slope of the calibration curves.

**Figure 3.**
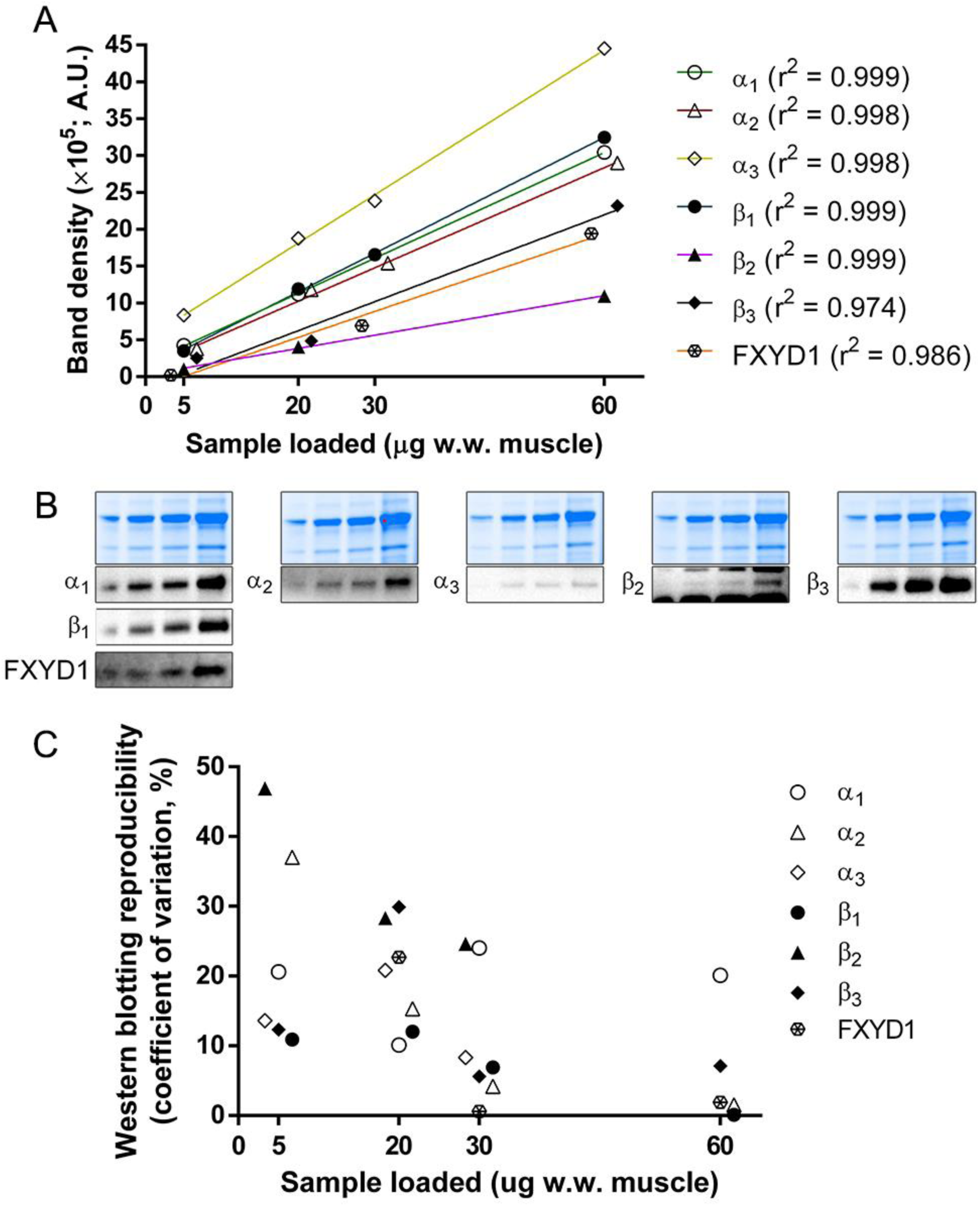
Representative calibration curves (A), blots (B), and western blotting reproducibility (C) for Na^+^,K^+^-ATPase isoforms and FXYD1. For every calibration curve, 5, 20, 30 and 60 μg w.w. of human vastus lateralis muscle was loaded onto the gels. To enable visualisation of all symbols, some data series in (A) and points in (B) were shifted by 1.70 units on the horizontal axis. Reproducibility was expressed as the coefficient of variation (CV), and calculated by normalising the density for a given loading amount to that of the slopes of the calibration curves for isoform blots and total protein on Stain Free gel. In B), total protein on Stain Free gels (*top panel*) and representative blots for the four-point calibration curves (*bottom panels*). Note that CV could not be calculated for the α_3_ and β_2_ isoforms for 60 μg w.w. muscle, as this point was excluded for some of the calibration curves due to saturation (cf. *Methods*).

### Statistical analysis

Statistical analyses were performed in Sigma Plot (Version 11.0, Systat Software, CA). Data was assessed for normality using the Shapiro-Wilk test. An appropriate transformation was used, if required, to ensure a normal distribution of data before subsequent analysis. A two-way repeated-measures (RM) ANOVA was used to evaluate the effects of time (Pre, Post) and fibre type (I and II) within group and using data from both groups (pooled data). A one-way ANOVA was used to assess the effect of group (COLD vs. CON) on Pre to Post change in protein content within fibre type (I and II). Data normalised to total protein, and not relative changes, were used for these analyses. Where applicable, multiple pairwise, *post hoc*, comparisons used the Tukey test. Cohen’s conventions were adopted for interpretation of effect size (*d*), where <0.2, 0.2-0.5, >0.5-0.8 and >0.8 were considered as trivial, small, moderate and large, respectively (Cohen, 1988). Data are reported as geometric mean ± 95% confidence intervals (CI95) in figures. Note that the protein expression at Pre is not equal to 1.0 due to the nature of using geometric means. F statistic (F), and *d* +/- CI95 are shown for time and fibre-type effects, as well as group interactions. The α-level was set at p ≤ 0.05.

## Results

In figures, individual data are displayed on the left with each symbol representing the same participant for all isoforms. On the right, geometric means ± CI95 are shown. Fold-changes are reported relative to participants’ geometric mean at Pre in type I fibres in CON.

### Validation of antibodies for immunoblotting

The results from our antibody validation are shown in Fig. 2.

The isoform specificity of the NKA α_1_ monoclonal antibody (#a6F) was verified using rat kidney and brain as positive controls (Fig. 2A). Our results support previous findings in similar tissues (Arystarkhova & Sweadner, 1996). We were also able to replicate the muscle type-specific distribution of α_1_ observed previously in rat skeletal muscles using other α_1_ antibodies (Lucchesi & Sweadner, 1991; Fowles *et al.*, 2004).

Specificity of the NKA α_2_ polyclonal antibody (#07-674) was verified by a clear band at the predicted molecular weight (105 kDa) in human and rat skeletal muscles, rat cardiac muscle and brain (Fig. 2B). These findings support previous results in the same tissues (Lucchesi & Sweadner, 1991; Thompson & McDonough, 1996; Wyckelsma *et al.*, 2015). We found that α_2_ was absent in rat kidney, which also corroborates with the literature (Hundal *et al.*, 1994; Lavoie *et al.*, 1996; Crambert *et al.*, 2002). BLAST analysis of the peptide sequence specific to the α_2_ antibody (#07-674, lot #2444088; aa sequence 432-445, human) revealed no cross reactivity with other NKA isoforms.

NKA α_3_ protein is highly expressed in rat brain (Lavoie *et al.*, 1997), but absent in rat cardiac muscle (Lucchesi & Sweadner, 1991; Sweadner *et al.*, 1994) and kidney (Hundal *et al.*, 1994). One study has shown that α_3_ may be absent also in rat skeletal muscle (Lavoie *et al.*, 1997). By use of the monoclonal NKA α_3_ antibody (#MA3-915), we support these previous findings (Fig. 2C). The multiple bands in human skeletal muscle could indicate multiple splice variants, as observed previously (Tumlin *et al.*, 1994).

The monoclonal NKA β_1_ antibody (#MA3-930) has been used previously in this journal to quantify β_1_ protein in human muscle samples despite no clear evaluation of specificity of the lot used (Murphy *et al.*, 2004). As β_1_ protein is heavily glycosylated (Vagin *et al.*, 2006), and highly expressed in human cancer cell lines (Salyer *et al.*, 2013), we tested the specificity of our β_1_ antibody lot using deglycosylated (PNGase treated for 3 h) and control human crude skeletal muscle samples and two human breast cancer (MDA-MB-231 and MCF10.Ca1d) and control (HEKs, human embryonic kidney) cell lines (Fig. 2D). Our findings of a substantial downshift of the predicted β_1_ band in deglycosylated samples and the markedly higher density of the same band in cancer cell lines vs. HEKs strongly support the specificity of our antibody lot for β_1_ protein. BLAST analysis of the peptide sequence specific to the β_1_ antibody (#MA3-930; aa sequence 195-199, sheep) confirmed absence of cross reactivity with other NKA isoforms.

Specificity of the polyclonal NKA β_2_ antibody (#22338-1-AP) was supported by detection of recombinant β_2_ protein (#Ag17818, ProteinTech; Fig. 2E, bottom panel). In further support, our antibody was able to detect a band at the predicted molecular weight in our positive (rat brain), but not in our negative (rat cardiac muscle), control sample. It also revealed a muscle type-specific expression of β2 in rat (Fig. 2E), in line with what has been found previously (Fowles *et al.*, 2004; Kristensen & Juel, 2010b). BLAST analysis of the peptide sequence specific to the β2 antibody lot used in the present study revealed that it did not cross react with other NKA isoforms.

Using the monoclonal NKA β_3_ antibody (#610992), we detected a band at the predicted molecular weight in human muscle (albeit weak in our whole-muscle, crude homogenate), rat soleus and cardiac muscle and kidney (Fig. 2F). Presence of β3 protein in rat kidney is in coherence with presence of β_3_ gene transcripts in the same tissue (Malik *et al.*, 1996). We found that β_3_ protein was absent in rat EDL muscle and brain, suggesting tissue-specific expression at the protein level.

Specificity of the polyclonal FXYD1 antibody (#13721-1-AP) was supported by detection of a band at the predicted molecular weight for our positive controls (human and rat skeletal muscles, rat cardiac muscle and kidney; Fig. 2G). FXYD1 is either not expressed or very lowly expressed in central nervous tissue (Crambert *et al.*, 2002). In accordance, we did not detect it in rat brain. The higher density of the band in rat soleus vs. EDL muscle supports previous findings (Rasmussen *et al.*, 2008). The longer migration of FXYD1 in the human vs. rat tissues could indicate specie- or case-specific differences in post-translational modification, such as phosphorylation (Fuller *et al.*, 2009; Thomassen *et al.*, 2016). BLAST analysis of the peptide sequence specific to the FXYD1 antibody (sequence 1-92 encoded by BC032800) showed no signs of cross reactivity with NKA isoforms.

### Western blotting technical error

The points constituting calibration curves on gels were strongly correlated (r^2^ ≥ 0.98, *n* = 22 gels, Fig. 3A). Representative blots for these curves are shown in Fig. 3B. The reproducibility of western blotting for each of the NKA isoforms and FXYD1 is shown in Fig. 3C. The technical error for these isoforms were ~10-30 % for the protein loading amount used (1.5 fibre worth of protein), isoform-dependent, and for most isoforms inversely related to the amount of protein loaded on each gel (Fig. 3C).

### Representative blots and verification of fibre type of fibre pools

Representative blots for training-induced effects on protein abundance, and verification of fibre type of fibre pools are shown in Fig. 4. It is clear from these results that fibres were grouped correctly, confirmed by the clear difference in MHC-isoform expression between type I and II fibre pools.

**Figure 4.**
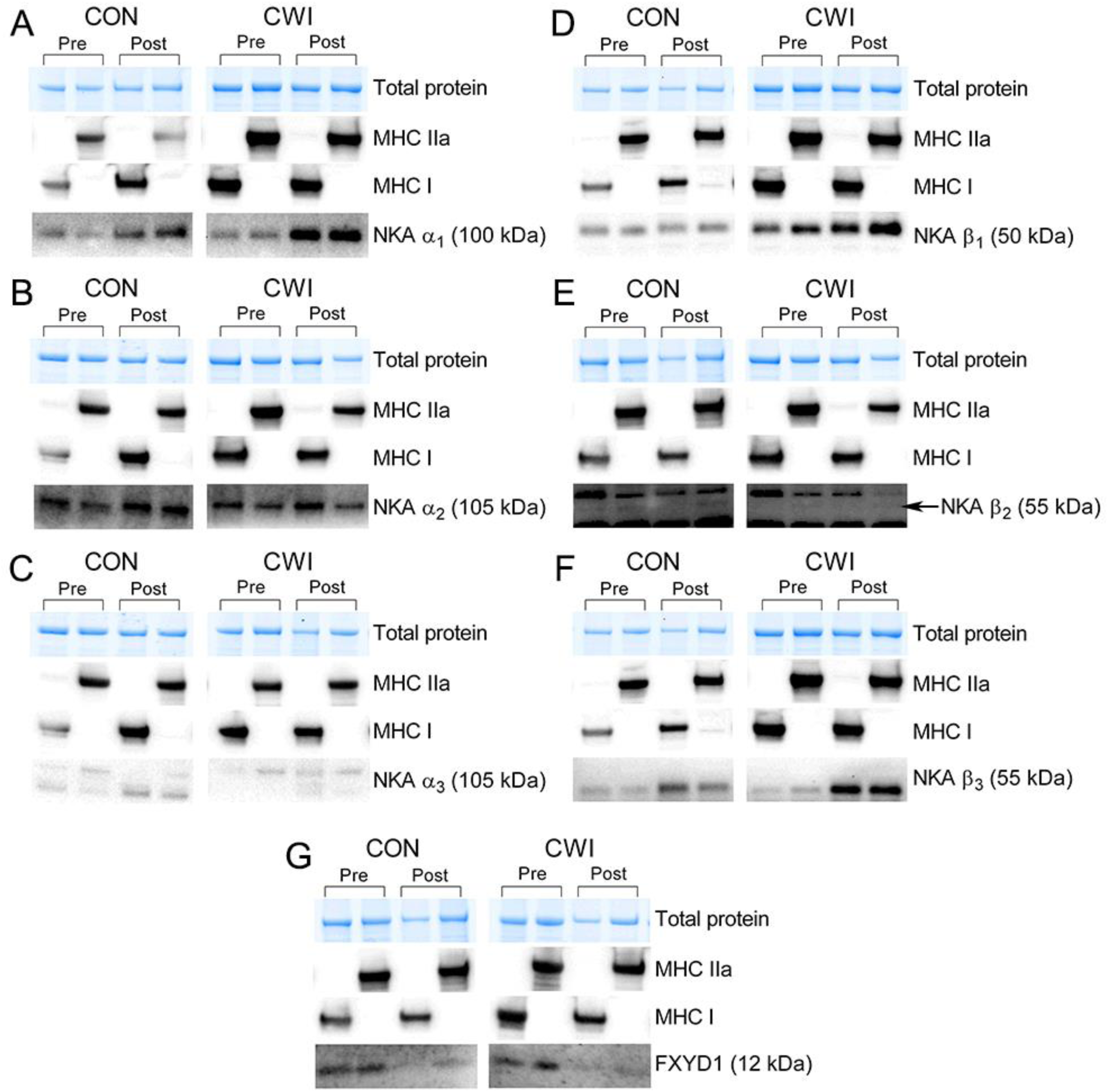
Representative blots for the effect of training without (CON) or with (CWI) cold-water immersion on NKA-isoform protein abundance. Total protein on Stain Free gels used for analysis (top panel), myosin heavy chain isoform expression of fibre pools (middle panel), and representative blots for NKA isoforms from the same run (bottom panel). Blots for NKA α_1_, α_2_ and α_3_ are shown in A, B and C, and NKA β_1_, β_2_, β_3_ and FXYD1 in D, E, F and G, respectively.

### Na^+^,K^+^-ATPase α_1_, α_2_, and α_3_

In CON, α_1_ protein increased with training (main effect for time: F = 19.78; p = 0.004; *d* = 1.41±0.77; *n* = 7) in both type I (p = 0.008; *d* = 1.20±0.80) and II fibres (p = 0.002; *d* = 1.56±0.78). Similarly, in COLD, α_1_ protein increased with training (main effect for time: F = 8.66; p = 0.042; *d* = 1.55±0.81; *n* = 4) in both type I (p = 0.053; *d* = 1.40±0.85) and II fibres (p = 0.029; *d* = 1.61±0.82). The increase in α_1_ protein was not different between groups in both fibre types (type I: p = 0.607; *d* = 0.07±0.04; type II: p = 0.348; *d* = 0.36±0.21; Fig. 5A). In both groups, α_2_ protein remained unchanged with training (main effect for time in CON: F = 3.65; p = 0.105; *d* = 0.34±0.33; *n* = 7; and in COLD: F = 0.69; p = 0.452; *d* = 0.48±0.34; *n* = 5; Fig. 5B). Based on the pooled data (both groups), α_2_ protein remained unchanged with training in both fibre types (type I: p = 0.161, *n = 12*; type II: p = 0.112, *n* = 12). Based on the same data, α_2_ protein was 17 ± 46 % more abundant in type II, compared to type I, fibres (main effect for fibre type: F = 9.63; p = 0.010; *d* = 0.38±0.26; *n* = 12). The α_3_ protein remained unchanged in both groups with training (main effect for time in CON: F = 3.71; p = 0.103; *d* = 0.53±0.52; *n* = 7; and in COLD: F = 0.55; p = 0.501; *d* = 0.03±1.02; *n* = 5; Fig. 5C).

**Figure 5.**
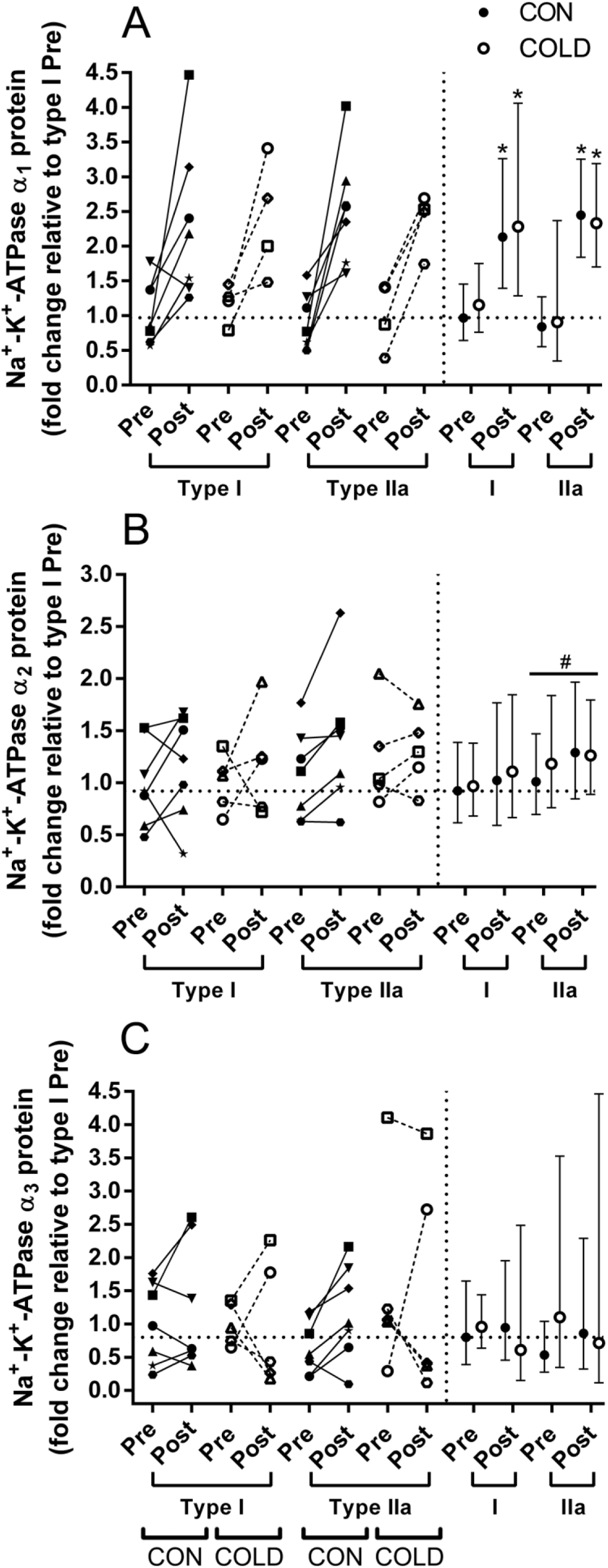
Effect of six weeks of repeated, intense training with (COLD) or without (CON) post-exercise cold-water immersion on Na^+^,K^+^-ATPase α-isoform protein abundance in type I and II human skeletal muscle fibres. A) α_1_, B) α_2_, and C) α_3_ protein abundance. Individual values (left) and geometric mean ± 95% confidence intervals (right) are displayed on each graph for CON (• closed symbols) and COLD (○ open symbols). Each symbol represents one participant (left) and is the same for protein and gene data (Fig. 2). The horizontal, dotted line represents the geometric mean expression at Pre in CON. Muscle was sampled at rest before (Pre) and after 6 weeks of training (Post). *p < 0.05, different from Pre within group; #p < 0.05, different from type I fibres based on pooled group data from both time points (please note the underscore of # for pooled group data).

### Na^+^,K^+^-ATPase β_1_, β_2_, and β_3_

β_1_ protein remained unchanged with training in both groups (main effect for time in CON: F = 1.25; p = 0.306; *d* = 0.37±0.45; *n* = 7; and in COLD: F = 0.01; p = 0.960; *d* = 0.10±0.58; *n* = 5; Fig. 6A). Based on the pooled data (both groups), β_1_ protein increased by 44 ± 75 % in type II fibres with training (p = 0.038; *d* = 0.53±0.35), but remained unchanged in type I fibres (p = 0.149; *d* = 0.29±0.24). Using the same data, β_1_ protein was 62 ± 80 % more abundant in type II than in type I fibres at Post (p = 0.003; *d* = 0.77±0.45, *n* = 12). As our western blots did not allow for quantitative assessment of training-induced effects on NKA β_2_ protein abundance, these data were excluded from analysis and are not presented. A representative blot is shown in Fig. 4E. Based on the pooled data (both groups), β_2_ protein was 54 ± 95 % more abundant in type II, compared to type I, fibres (main effect for fibre type: F = 14.84; p = 0.003; *d* = 0.56±0.56; *n* = 11). In CON, β_3_ protein increased with training (main effect for time: F = 38.62; p < 0.001; *d* = 1.23±1.05; *n* = 7) in both type I (p = 0.003; *d* = 1.18±0.97) and II fibres (p < 0.001; *d* = 1.24±1.15). Similarly, in COLD, β_3_ protein increased with training (main effect for time: F = 22.40; p = 0.009; *d* = 1.18±2.93; *n* = 5) in both type I (p = 0.006; *d* = 1.18±2.93) and II fibres (p = 0.012; *d* = 1.21±1.92). The increase in β_3_ protein was not different between groups in either fibre type (group × time interaction: F = 0.59, p = 0.459 for type I; and F = 0.04, p = 0.854 for type II). Based on the pooled data (both groups), β_3_ protein increased with training (main effect for time: F = 64.94; p < 0.001; *d* = 1.13±1.19; *n* = 12) in both type I (2.55 ± 2.39 fold; p < 0.001; *d* = 1.07±1.35) and II fibres (2.22 ± 1.81 fold; p < 0.001; *d* = 1.22±1.03; Fig. 6B).

**Figure 6.**
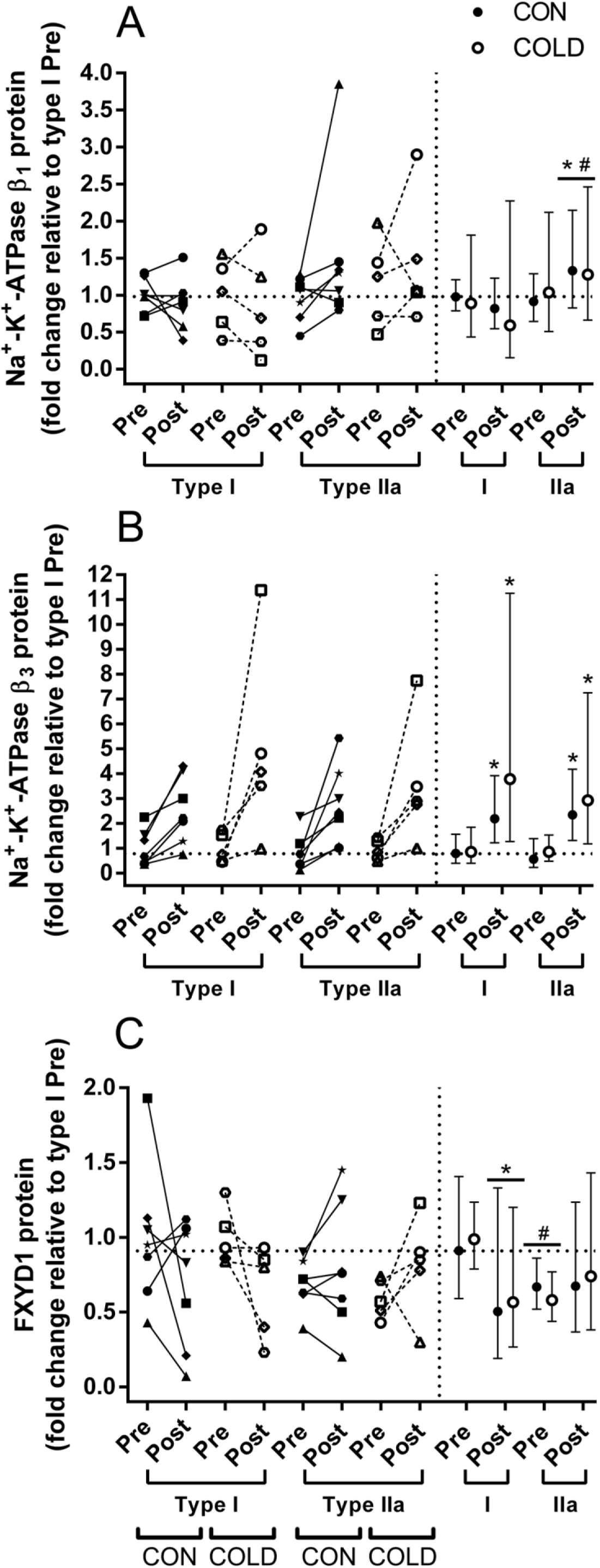
Effect of six weeks of repeated, intense training with (COLD) or without (CON) post-exercise cold-water immersion on Na^+^,K^+^-ATPase β-isoform and FXYD1 protein abundance in type I and II human skeletal muscle fibres. A) β_1_, B) β_3_, and C) FXYD1 protein abundance. Individual values (left) and geometric mean ± 95% confidence intervals (right) on each graph for CON (• closed symbols) and COLD (○ open symbols). Each symbol represents one participant (left) and is the same for protein and gene data (Fig. 3). The horizontal, dotted line represents the geometric mean expression at Pre in CON. Muscle was sampled at rest before (Pre) and after 6 weeks of training (Post). Note the different scale on the secondary axis and that NKA β2 protein data were excluded from analysis. *p < 0.05, different from Pre within group; *p < 0.05, different from pooled group data at Pre; #p < 0.05, Fibre type difference within time point; #p < 0.05, Fibre type difference within time point for pooled group data (please note the underscore of # and * for pooled group data).

### Phospholemman (FXYD1)

In both groups, FXYD1 protein also remained unchanged with training (main effect for time in CON: F = 1.56, p = 0.258, *d* = 0.66±0.34, *n* = 7; and in COLD: F = 0.23, p = 0.654, *d* = 1.17±0.27, *n* = 5). Based on the pooled data (both groups), FXYD1 protein decreased by 33 ± 40 % in type I (p = 0.012; *d* = 0.82±0.22), but remained unchanged in type II fibres (p = 0.535; *d* = 0.51±0.17). Based on the same data, FXYD1 protein was 35 ± 35 % more abundant in type I, compared to type II, fibres at Pre (main effect for fibre type: F = 6.31; p = 0.020; *d* = 1.01±0.57; *n* = 12), but not at Post (F = 0.79; p = 0.384; *d* = 0.31±0.17; *n* = 12; Fig. 6C).

## Discussion

The main novel findings, which are summarised in Fig. 7, were that six weeks of intense exercise training increased NKA α_1_ and β_3_ protein abundance in both fibre types and β_1_ protein in type II fibres, but had no impact on α_2_ and α_3_ abundance in either fibre type. Furthermore, training decreased FXYD1 protein content in type I fibres and abolished its fibre type-dependent expression detected before training. These results support that improvements in muscle K^+^ regulation after a period with repeated, intense exercise sessions in humans (McKenna, 1995; Nielsen *et al.*, 2004) may, in part, be attributable to fibre type-specific modulation of NKA-isoform protein abundance. This fibre type-dependent regulation of NKA could be one explanation for the dissociation between muscle NKA activity and whole-muscle NKA-isoform protein content previously reported after a period of intense training in humans (Aughey *et al.*, 2007). Furthermore, CWI neither adversely nor favourably affects training-induced adaptations in NKA-isoform abundance in different muscle fibre types. The technical error of western blotting for the NKA isoforms was ~10-30 % and isoform-dependent.

**Figure 7.**
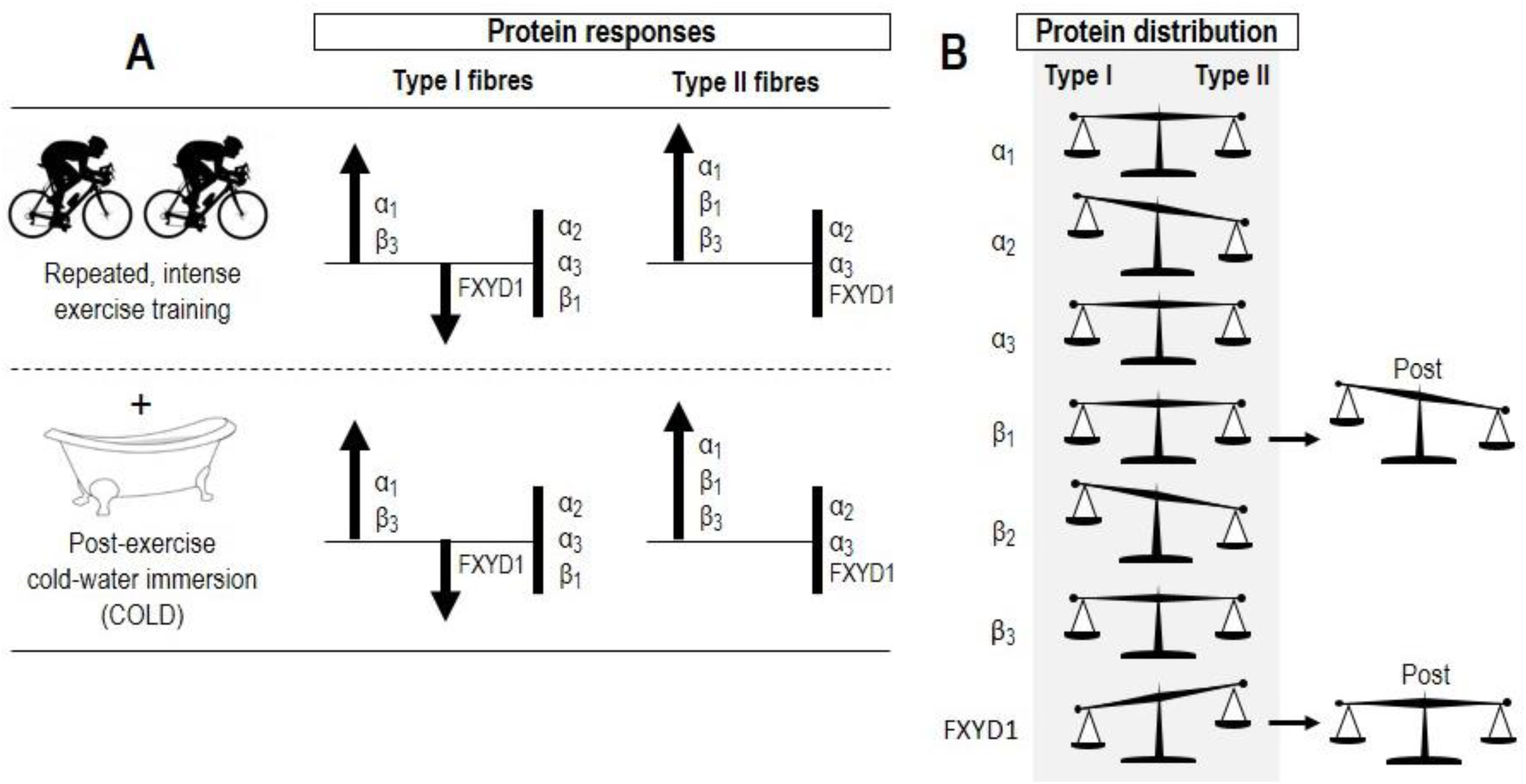
Summary of key findings. A) Effect of repeated, intense exercise training on the protein abundance of Na^+^,K^+^-ATPase isoforms (α_1-3_ and β_1-3_) and phospholemman (FXYD1) in different muscle fibre types (type I and type II). Bold vertical lines without arrow indicate NKA isoforms that remained unchanged with the given intervention. B) Protein distribution of Na^+^,K^+^-ATPase isoforms (α_1-3_ and β_1-3_) and phospholemman (FXYD1) in type I and type II fibres, and the effect of the training period on this distribution (Post).

### Regulation of NKA α isoforms in type I and II human muscle fibres by repeated-intense training

A novel finding of the present study was that NKA α_1_ protein abundance increased in both fibre types with training (>2 fold, Fig. 5A). This indicates that α_1_ protein content may be regulated similarly in type I and II muscle fibres in response to sprint interval training in humans. In contrast, training was without effect on α_2_ and α_3_ protein content. No effect of sprint training on α-isoform (α_1_, α_2_ and α_3_) abundance has previously been detected in isolated type I and II human skeletal muscle fibres (Wyckelsma *et al.*, 2015), although the duration of both the sprints (4 s) and the training period (4 weeks) was shorter. However, other human studies have reported increases in [^3^H]-ouabain binding site content, reflecting increased α-isoform content, with sprint interval training (McKenna *et al.*, 1993; Harmer *et al.*, 2006). Our present results suggest these increases in binding may, in part, be related to elevated α_1_ protein content in both fibre types. While lack of sufficient muscle precluded measurement of [^3^H]-ouabain binding site content, our results support that increases in α_1_ protein content may also be important for improvements in a muscle’s capacity for K^+^ regulation during repeated, intense exercise in humans (McKenna *et al.*, 1993).

Although there was not a significant increase in NKA α_2_ protein content after training (pooled data: p<0.07, *d* = 0.42), α_2_ protein content was quantitatively higher in around three quarters of individual type I and II fibre pools. This suggests a possible upregulation in α_2_ that was not detected due to inter-individual variability. Biological variability could perhaps also have influenced this result given the low number of fibre segments (range: 2-16) contained in each fibre pool. However, this explanation seems less likely, since AMPKβ_2_, SERCA1 and actin can be validly quantified in pools consisting of 2-20 fibre segments, and pools of few segments (*n* = 4) are reproducible for quantitative protein analysis (unpublished observation). In addition, consistent with a previous observation in recreationally active men (37 %; Thomassen *et al.*, 2013), we found higher α_2_ protein content in type II vs. type I fibres (~17 %; Fig. 8B). This raises the possibility that the ion transport function of NKA may be fibre type-dependent in humans, in line with what has been shown in rats (Kristensen & Juel, 2010a). No fibre-type difference for α_2_ protein content was detected in subjects with a substantially lower VO_2peak_ relative to our participants (Wyckelsma *et al.*, 2015). Thus, in humans, higher training status may be associated with greater type-II muscle fibre α_2_ protein content.

Little is known about the relevance of NKA α_3_ in skeletal muscles. Consistent with our current finding, the α_3_ protein content remained unchanged at the whole-muscle level after 3 weeks of cycling (8 × 5-min at 85% VO_2max_) in well-trained men (Aughey *et al.*, 2007). Albeit not quantified in absolute terms, our validation blots for α_3_ revealed that this isoform is lowly expressed at the protein level in human skeletal muscle (Fig. 2). This, along with the low α_3_ mRNA expression detected previously in the same tissue (Aughey *et al.*, 2007), downplays the functional importance of α_3_ for the muscle’s contractile function in humans.

### Regulation of NKA β isoforms in type I and II human muscle fibres by repeated-intense training

In a previous human study, four weeks of repeated-sprint training selectively increased the protein content of NKA β_1_ in type II muscle fibres (identified in individual fibre segments) (Wyckelsma *et al.*, 2015). In agreement, in the present study, β_1_ protein abundance was higher (44%) after training in type II fibres only. These observations, along with the increase in α_1_ abundance in the present study, could indicate a need for improved NKA activity in this fibre type with intense training. In support, in rat gastrocnemius muscle, higher NKA hydrolytic activity was reported in membrane vesicles with a reduced (50 %) molar α_2_/β_1_ ratio caused by higher β_1_ content, relative to vesicles with a greater ratio (1.0) (Lavoie *et al.*, 1997). Although our data did not allow for quantification of the possible impact of training on β_2_ protein content, we were able to confidently show that β_2_ protein abundance is higher in type II, relative to type I, fibres (Fig. 4E). This is in accordance with a previous finding in individual fibre segments (27 % higher in type II fibres) (Wyckelsma *et al.*, 2015). As the K_m_ for Na^+^ of α/β_2_ heterodimers (7.5-13 mM) is higher than the corresponding K_m_ for α/β_1_ complexes (4-5.5 mM) in rat skeletal muscles (Kristensen & Juel, 2010a), our findings raise the possibility that human muscle NKA activity may also be fibre type-dependent. Further research is needed to determine if NKA activity is fibre type-dependent in human muscle. It has been reported in humans that NKA β_3_ protein content is substantially elevated with age in whole muscle (2.5 fold), and in type I (1 fold) and II (3 fold) skeletal muscle fibres (McKenna *et al.*, 2012; Wyckelsma *et al.*, 2016). In rat skeletal muscles, a similar age-associated rise in β_3_ content was reversed by ~14 weeks of endurance training (Ng *et al.*, 2003), indicating that regular, continuous muscle activity potently affects β_3_ content with age. We demonstrate here that brief, repeated-intense training substantially increases NKA β_3_ protein content (increase > 2 fold) in type I and II human muscle fibres. Thus, the type of muscle activity seems critical for β_3_ content of both muscle fibre types in humans. The rise in β_3_ content occurred in parallel with elevated α_1_ protein abundance (Fig. 5A and 6B), suggesting an enhanced potential for α_1_/β_3_ complex assembly in both fibre types post training. This supports that the β_3_ isoform could take part in the maintenance of resting membrane potential in both contracting and non-contracting muscle fibres, in line with the ion transport function of α_1_ (Radzyukevich *et al.*, 2013), although it could exert other yet unidentified functions in skeletal muscles.

### Regulation of phospholemman (FXYD1) in human muscle fibre types by repeated, intense training

Previous human studies using whole-muscle samples reported no alterations in FXYD1 protein abundance after 10 days to 8 weeks of intense training (Thomassen *et al.*, 2010; Benziane *et al.*, 2011; Nordsborg *et al.*, 2012; Skovgaard *et al.*, 2014). Conversely, we found that six weeks of intense training decreased FXYD1 abundance by 33 % in type I fibres (Fig. 6C). This decrease is reinforced by the large effect size (0.82), small confidence interval (22 %), and good reproducibility (CV < 22 %; Fig. 3C). However, FXYD1 abundance was unchanged in type II fibres (Fig. 6C), highlighting for the first time that FXYD1 is regulated in a fibre type-dependent manner by intense training in human muscle. These results underline that physiologically relevant adaptations may have been overlooked in the previous human studies due to their fibre type heterogeneous samples. But their use of sample fractionation, i.e. removal of an indefinite amount of protein (Murphy & Lamb, 2013), may also have influenced these previous outcomes. Co-localisation of unphosphorylated FXYD1 with α/β heterodimers may inhibit their activation by increasing K_m_ (i.e. decreased affinity) for Na^+^ and K^+^ (Crambert *et al.*, 2002), whereas the interaction of FXYD1 with α_1_ or α_2_ isoforms remains unaffected by exercise (Benziane *et al.*, 2011). This suggests that the decline in type-I fibre FXYD1 abundance in the current study, independent of contraction-stimulated effects on its co-localisation with NKA α/β complexes *per se,* may have been functionally important. Future training studies should combine coimmunoprecipitation analyses and measures of NKA function (e.g. maximal *in vitro* activity or muscle K^+^ release during exercise), with fibre type-specific protein analyses to evaluate this possibility.

### No effect of CWI on the effect of training on NKA-isoform protein abundance in different human muscle fibre types

In the present study, post-exercise CWI was without effect on the fibre type-specific changes in NKA-isoform protein abundance with training. This underlines that CWI can be performed regularly after intense training sessions (or match play) by athletes, to utilise its potent placebo effect (Broatch *et al.*, 2014), without adversely affecting training-induced changes in NKA-isoform abundance, and possibly, NKA function. Whether an absence of a CWI effect relates to an insufficiency to activate the molecular signalling events prerequisite to transcriptional and/or post-transcriptional modification cannot be resolved from the present study. However, our unpublished data reveal that a single session of post-exercise CWI transiently increases NKA α_2_ mRNA content in skeletal muscle of healthy men, indicating a bout of CWI may augment the transcription, stabilisation, or both, of this NKA mRNA transcript in human muscle. The present results highlight that this transient effect of CWI does not translate into increased protein abundance after weeks of training. This raises the possibility that α_2_ protein content is regulated, in part, by factors other than an increased potential for mRNA translation induced by higher mRNA content in response to repeated exercise sessions in human muscle.

### Reliability of western blotting for NKA isoforms

In many human studies, inferences about training-induced effects on the capacity of skeletal muscle for ion regulation were based, in part or fully, on modest changes (9-39 %) in protein abundance quantified using western blotting (Green *et al.*, 2004; Nielsen *et al.*, 2004; Iaia *et al.*, 2008; Thomassen *et al.*, 2010; Gunnarsson *et al.*, 2012). This is surprising considering these studies were limited by the use of fibre-type heterogeneous, fractionated samples (Murphy & Lamb, 2013), and, for some, no consideration of blot linearity (Mollica *et al.*, 2009) nor antibody lot validation. With the use of Stain Free imaging technology, wet transfer, and highly-sensitive chemiluminescence, we report that in our hands the technical error of western blotting using protein from a fibre segment of ~1-3 mm in length, amounting to ~15 μg wet weight tissue, is of a similar magnitude (~10-30 %) as some of these previously reported changes (Fig. 3C). Our results highlight the importance of taking into account the reliability of western blotting when interpreting changes in muscle NKA-isoform protein abundance.

### Conclusions and perspectives

The effectiveness of intense intermittent training to improve the capacity of skeletal muscle for transmembrane Na^+^/K^+^ transport in humans is well-documented. The present study revealed that these improvements may be ascribed, in part, to selective modulation of the protein abundance of NKA β_1_ in type II fibres, FXYD1 in type I fibres, and NKA α_1_ and β_3_ in both fibre types. The insights gained from this study improve our understanding of how NKA function may regulated in different muscle fibre types to accommodate the need for Na^+^ and K^+^ transport during intense, intermittent exercise in human skeletal muscle. Furthermore, our data highlight that CWI can be performed on a regular basis without adversely affecting training-induced modulation of NKA-isoform protein abundance. Future work should assess protein responses in different muscle fibre types in combination with measurements of muscle K^+^ handling to elucidate the functional relevance of these fibre type-dependent protein responses to training.

## Additional information

## Author contributions

Exercise testing and training were performed at Institute of Sport, Exercise and Active Living (ISEAL), Victoria University, Melbourne, VIC 3011. Antibody validation, reproducibility, and protein analyses were performed at the Department of Biochemistry and Genetics, La Trobe University, Melbourne, VIC 3086, Australia. All authors contributed to drafting and critically revising of this manuscript. DC, RMM, JRB and DJB contributed to the conception, design of experiments, and collection and analysis of data. DC RMM, JRB, MJM and DJB contributed to data interpretation. All authors approved the final version of this manuscript.

## Acknowledgements

We thank our participants for kindly donating parts of their muscle and for their efforts throughout the study. The monoclonal antibodies used to measure myosin heavy chain isoforms and the NKA α_1_ isoform were obtained from the Development Studies Hybridoma Bank (DSHB) under the auspices of the NICHD and maintained by the University of Iowa, Department of Biological Sciences, Iowa City, IA 52242, USA. The antibodies directed against adult human MHC I and IIa isoforms (A4.840 and A4.74, respectively) were developed by Dr H. Blau, and the antibody directed against NKA α_1_ (a6F) was developed by Dr D.M. Fambrough. We thank Hendrika Duivenvoorden, LaTrobe University, for providing the breast cancer cell lines.

